# A test of limited transpiration traits in sorghum to improve late-season water use, photosynthesis, growth, and grain yield in the high plains of Northern Colorado

**DOI:** 10.1101/2024.06.27.601079

**Authors:** Sean M. Gleason, Jared J. Stewart, Stephanie K. Polutchko, Brendan S. Allen

**Affiliations:** Water Management and Systems Research Unit, USDA-ARS, Fort Collins, CO 80526, USA

**Keywords:** Limited transpiration, staygreen, crops, water use, water conservation

## Abstract

Limited transpiration (LT) traits aim to conserve early-season water to benefit late-season grain development. While theoretical and modeling efforts support LT efficacy, empirical tests directly measuring water loss from leaves and canopies are scarce. This study evaluates the performance of LT genotypes in achieving reduced early-season water use and improved late-season growth and yield in semi-arid Colorado. The research involved near-isogenic lines (NILs) derived from sorghum inbred lines, subjected to different irrigation treatments. Measurements included stomatal conductance, net CO_2_ assimilation, and photosystem II (PSII) efficiency. Results indicate that LT genotypes did not consistently exhibit lower early-season water use or higher late-season growth compared to non-LT genotypes. Early-season water use was *positively* correlated with above-ground biomass, challenging the assumption that early-season water conservation can be leveraged for late-season benefits. We question the efficacy of LT traits, highlighting the physiological link between water use and carbon gain, and the potential opportunity costs of reduced early-season growth. We suggests that breeding strategies should focus on enhancing deep soil water access and maximizing carbon gain rather than merely reducing transpiration or shifting water use in arid and semi-arid environments.

## Introduction

Limited transpiration traits (hereafter “LT”) include leaf level, canopy level, and whole-plant level traits that serve to reduce early-season water use such that water can be stored in the soil without meaningful loss until it is needed for late-season grain development (Sinclair 2018). The theory underpinning LT traits is over two decades in the making, and has broad support from funding agencies, crop physiologists, and seed companies (Sinclair et al. 2017). Although modeling efforts support the theoretical efficacy of LT traits in some soil and climate contexts (Sinclair et al. 2005; Hammer et al. 2014; Messina et al. 2015; Gleason et al. 2022), surprisingly, there are almost no empirical tests of this theory that include direct measurements of water loss from leaves and canopies, despite claims that these processes underpin LT trait efficacy – i.e., reduced early-season stomatal conductance, transpiration, gas-exchange. Rather, previous field studies have relied almost entirely on soil water flux, grain yield, and other non-physiological traits that are often measured in breeding programs (e.g., plant height, ear and grain characteristics, “greenness” indices) (Reyes et al. 2015; Sinclair 2018). Although there has been much published on LT traits and genotypes, the vast majority of this literature has focused on specific traits/mechanisms indirectly related to LT (e.g., “greenness”, timing of leaf senescence) and the genetic underpinnings of these traits (e.g., Borrell et al. 2000; Gholipoor et al. 2010, 2013; Wang et al. 2012; Choudhary and Sinclair 2014; Shekoofa et al. 2014), without testing the prediction that “LT” genotypes exhibit reduced early-season water use and higher grain yield than non-LT genotypes. As such, in our view, current research on LT traits and genotypes lack a sound empirical foundation for their investment.

There are at least two theoretical reasons why we should not expect LT traits to result in improved growth and yield in at least some cases. The first reason arises from the positive causal relationship between stomatal conductance (g_S_) and net CO_2_ assimilation (A_net_), i.e., plants must “spend” water to obtain atmospheric CO_2_. Although the A_net_ ∼ g_S_ relationship is near-proportional between 0 and 0.3 mol m^-2^ s^-1^ in most crop species, the slope of this relationship becomes shallower beyond this point and plants must spend more water to obtain the same CO_2_ increment. Much has been made about the importance of this diminishing return on water expenditure, especially in C3 species which lack Kranz anatomy and the accompanying internal CO_2_-concentrating biochemistry. However, given that C4 species, like sorghum, typically exhibit monotonic increasing A_net_ ∼ g_S_ functions, rather than saturating functions, there exists less opportunity for C4 species (and likely C3 species) to save water by limiting maximal stomatal conductance. This is because vascular species “self-regulate” both gas-phase (Buckley 2015, 2017) and liquid-phase (Scoffoni et al. 2011; Ocheltree et al. 2020) conductances at low water potential, thus limiting their water use, even in the absence of LT traits. The second reason why we may not expect LT traits to confer improved growth and yield arises from the consequences of forgoing carbon gain early in the season. Atmospheric carbon that is not fixed by the plant early in the season cannot be invested in structural (e.g., roots, leaves, stems) and non-structural carbohydrates. Importantly, given the exponential nature of resource investment on plant growth, the earlier water (and carbon) reductions are implemented, the more costly it will be for the plant (Hunt 1978; Holland et al. 2019). The second consequence of “saving” water in the soil for up to several months is that it cannot *all* be saved for late-season transpiration, and thus, some fraction of this “banked” soil water will be lost to competing sinks (e.g., evaporation, saturated and unsaturated flow out of the rhizosphere, uptake by weeds), rather than passing through crop stomata. Importantly, the magnitude of these soil water losses are strongly climate- and soil-dependent (Monteith and Unsworth 1990; Allen et al. 2005; Passioura 2006), e.g., evaporative losses are expected to range between 20 and 87% of total crop evapotranspiration, depending on soil and atmospheric conditions (Fulton, J.M. 1966; Allen et al. 2005; Logsdon et al. 2014; Unkovich et al. 2018). Taken together, the monotonic increasing relationship between net CO_2_ assimilation and stomatal conductance, the potential opportunity costs associated with forgoing early season carbon capture, and competing alternative sinks for soil water (other than crop transpiration), give ample cause to question the efficacy of LT traits.

We evaluated the capacity of LT inbred lines to achieve lower early-season water use and improved late-season growth and yield in the semi-arid high plains of Colorado. Specifically, we quantified the efficacy of LT genotypes to: 1) achieve reduced water use during early season (pre-flowering) relative to non-LT genotypes, and 2) achieve higher rates of gas exchange (water vapor and carbon dioxide) during late season (post flowering), resulting from in-soil water savings during the pre-flowering period. 3) achieve higher late-season above-ground biomass and grain yield, relative to non-LT genotypes, via *lower* early-season water use. We address these questions at both leaf-specific and “whole-plant” levels.

## Methods

### Site, growth conditions, and weather

Plants were grown at the USDA Limited Irrigation Research Farm (LIRF) in the semi-arid climate of north-central Colorado during the 2021 and 2022 growing seasons. The soils at the site consist of deep clay loam Ustic Haplargids. Soil testing was carried out prior to planting each year and fertilization (N, P, K, Fe) was based on regional recommendations for sorghum. Pre-emergent herbicide was applied 14 days prior to planting and subsequent weeding was performed by hand. Water was applied via a variable rate irrigation linear sprinkler system (Lindsay Zimmatic, Omaha, NE, USA). Planting population was 185,000 seeds ha^-1^ in 2021 and either 200,000 seeds ha^-1^ (high) or 100,000 seeds ha^-1^ (low) in 2022. Grain yield results for 2022 reported here are pooled across both population treatments because there were no significant population effects on grain yield.

“Early-season” and “late-season” periods were determined as the time intervals between planting and flowering (first genotype to exhibit flowers), or between flowering and harvest, respectively. For example, in 2021, planting occurred on June 9 and flowering was first observed on August 25; therefore, the early-season period in 2021 was calculated as the time between June 9 and August 25 for all genotypes. Harvest occurred in 2021 on November 28. Planting, flowering, and harvest in 2022 occurred on May 26, August 14, and November 21, respectively. Daily precipitation, air temperature, and the vapor pressure deficit of the atmosphere are reported in the supplemental materials (SI Fig. 1). Total cumulative early-season and late-season precipitation in 2021 was 94 mm and 22 mm, respectively. In 2022, total cumulative early-season and late-season precipitation was 75 and 41 mm, respectively (SI Fig. 1).

### Treatments and genotypes

Plants were subjected to one of three water treatments: 1) fully-watered during the entire field season (hereafter “full-water”) such that 100% of the crop’s transpiration demands were satisfied, 2) early-season stress, where no irrigation water was applied between planting and flowering time, but plants were fully-watered thereafter, and 3) late-season stress, where plants were given full water between planting and flowering dates but were given no water thereafter.

Genotypes chosen for this study included near-isogenic lines (NILs) derived from inbred lines BTx642 and RTx7000. The F1 plants from this cross were backcrossed with RTx7000 to create NILs containing only one (of four) staygreen loci from BTx642 (Harris et al. 2007), i.e., SG1, SG2, SG3, or SG4, which had been identified in previous studies (Tuinstra et al. 1997; Crasta et al. 1999; Tao et al. 2000; Xu et al. 2000). Previous studies have reported that these staygreen loci are associated with reduced tillering, reduced leaf size, reduced canopy size at anthesis, reduced whole-canopy transpiration, and reduced stomatal density, relative to inbred line RTx7000 (Borrell et al. 2014a, 2014b). Non-LT genotypes (DKS54-00 and Tx430) were also planted and were compared against the LT genotypes. Hereafter, genotypes in the text and figures are referred to as SG1, SG2, SG3, SG4 (see above), and BTX (BTx642), whereas non-LT genotypes are referred to as TX (Tx430) or FILL (DKS54-00).

### Plant measurements

plant water use was measured using gas exchange systems (Licor 6400 equipped with a 6400-40 PAM chlorophyll fluorometer) and steady-state porometers (Decagon SC-1). Leaf-level water use and photosynthesis were scaled up to whole-plants by multiplying leaf-level measurements by the existing green leaf area existing at the time of measurement. To do this, green and senesced leaf area was measured at 43 d, 63 d, 79 d, and 106 d after planting. Green leaf area was estimated on measurement days that fell between these dates by fitting spline functions to green leaf area vs time for each genotype-by-treatment combination, and interpolating green leaf area from these functions.

Stomatal conductance was measured using three steady-state leaf porometers (Decagon SC-1) and two gas-exchange systems (Licor-6400). Stomatal conductance was measured using the Decagon SC-1 across eight early-season days in July (7/20 - 7/22, 7/26 – 7/30). Stomatal conductance, maximal net CO_2_ assimilation, photosystem II (PSII) efficiency, and PSII thermal dissipation were measured using a Licor-6400 equipped with a 6400-40 fluorometer across three early-season days (8/03 – 8/05), as well as two late-season days (9/13, 9/14). In total, 1389 measurements were taken across 13 days in July, August, and September.

### Statistical analyses

Daily trajectories of stomatal conductance were fit with quadratic models using the nlsLM and nlmer functions in the minpack.lm and lme4 statstical packages for R, respectively. Mixed-effects models were fit as a split-plot design using nlmer, with the three fitted quadratic coefficients, genotype, and treatment as fixed factors, and genotype within block as a random factor. All figures and analyses were done R version 4.1.2.

## Results

### Hypothesis 1: Staygreen genotypes exhibit less leaf-specific water use during early season than non-staygreen genotypes

BTX had lower mean and median stomatal conductance, net CO_2_ assimilation, and PSII efficiency during early-season measurements than non-staygreen genotypes, although only one of these comparisons was significantly different (PSII efficiency: BTX = 0.210 +/- 0.035, FILL = 0.243 +/- 0.032, p=0.0254) (Fig. 1). Late-season (post-flowering) stomatal conductance, net CO_2_ assimilation, PSII efficiency, and PSII thermal dissipation were similar among all genotypes during the late-season (Fig. 1C). Daily trajectories of stomatal conductance during the early season were similar across all genotypes (Fig. 2). However, the lower stomatal conductance exhibited by BTX during early season measurements (Fig. 1A), although not statistically significant, arose from lower stomatal conductance later in the day (ca. after 11am) (Fig. 2, “Pre-flower, full water”).

**Fig. 1.**
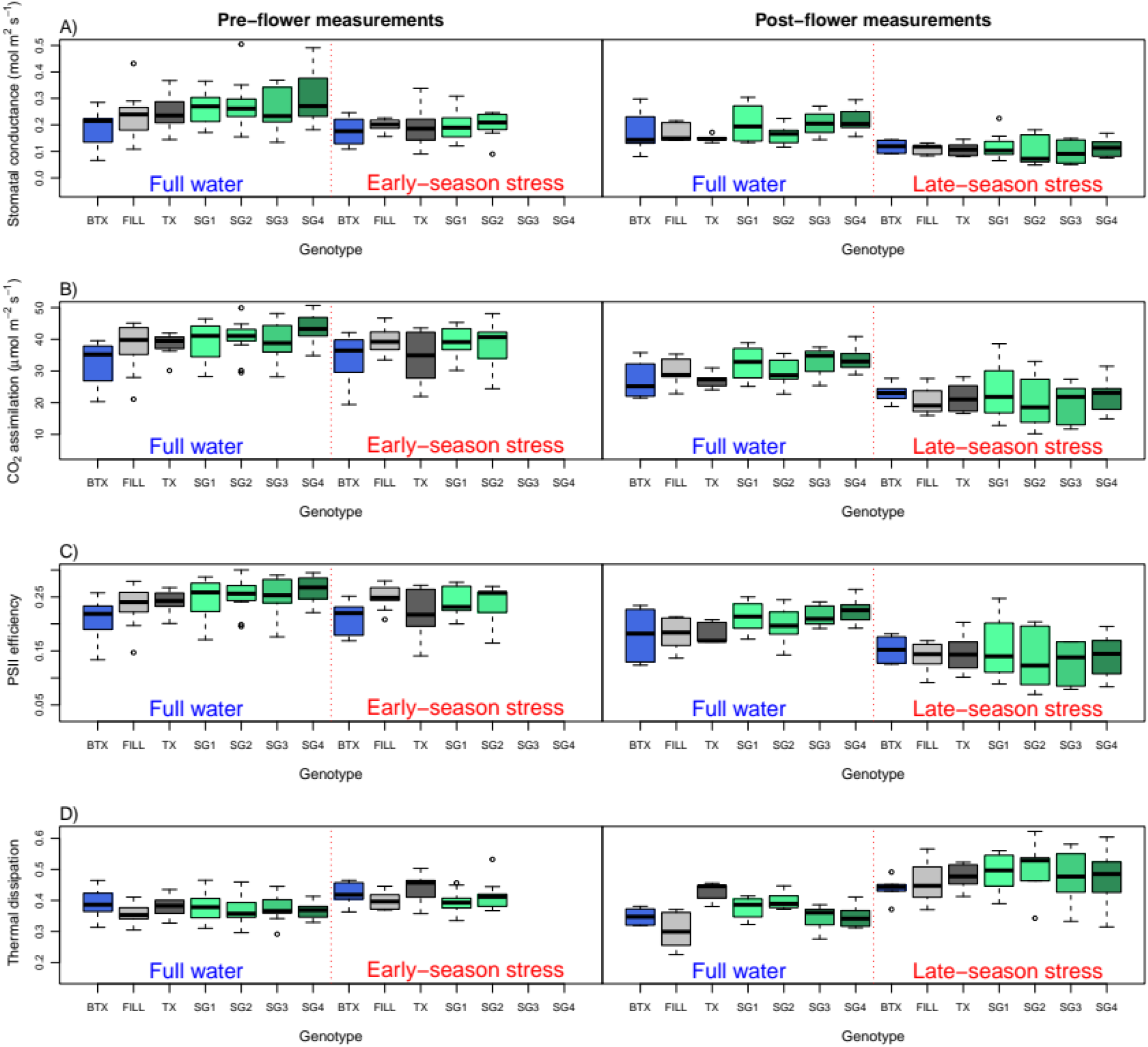
Stomatal conductance, net CO_2_ assimilation, and photosystem II efficiency and thermal dissipation for all genotypes across all treatments and time periods. All measurements were taken using a Licor 6400xt gas exchange systems equipped with 6400-40 PAM chlorophyll fluorometer.

**Fig. 2.**
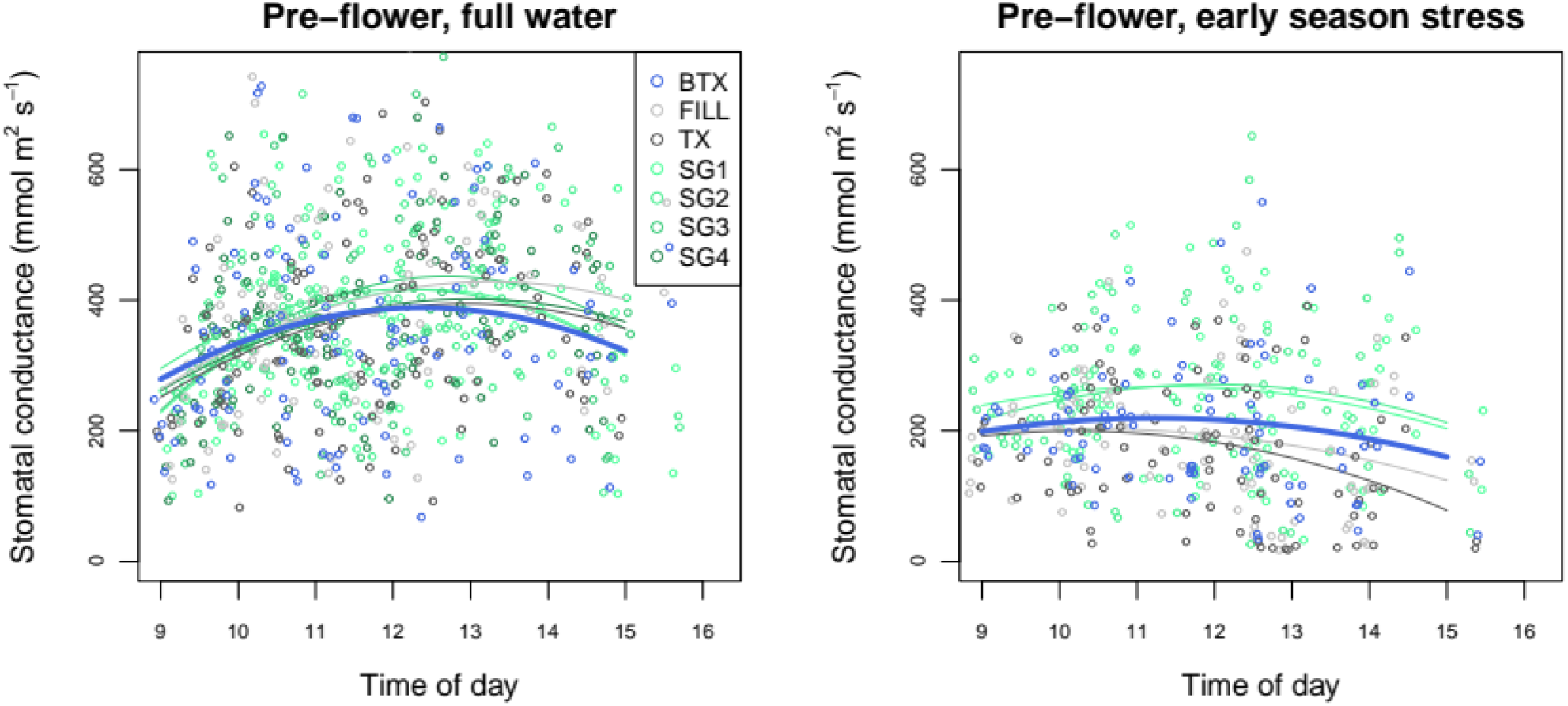
Daily time courses of stomatal conductance measured using a steady-state leaf porometer (Decagon SC-1) across eight days in July, 2022 (07/20 - 07-30). Data are fit with quadratic models. BTX is denoted with dark blue symbols and wide trendline.

### Hypothesis 2: Staygreen genotypes achieve higher rates of leaf-specific gas exchange (water vapor and carbon dioxide) during late season (post flowering), resulting from in-soil water savings during the pre-flowering period

Genotypes achieving the highest rates of gas exchange and stomatal conductance during the post-flowering period did not exhibit less water use during the pre-flowering period (Fig. 1). More generally, there was no negative correlation across genotypes between water use during the post-flowering period vs water use during the pre-flowering period. As such, the water use data do not support the assumption that lower rates of water use during the pre-flowering period can be used, or leveraged to greater effect, during the post-flowering period, under the soil and climate conditions considered here, at least when evaluated at the leaf level, i.e., not taking into account whole-plant transpiration and leaf loss during the season.

### Hypothesis 3: Staygreen genotypes achieve higher late-season biomass, relative to non-staygreen genotypes, via lower early-season water use

Similar to the pre-to-post flowering water use trends presented above, late-season (79-day) above-ground biomass was not inversely correlated with early-season water use, either at the leaf-level or whole-plant levels. Rather, whole-season growth and above-ground biomass accumulation by day 79 was strongly and *positively* correlated with early-season water use (Fig. 3). We do not present our analyses of grain yield here because many of the plots were not harvestable at the end of the season, however, we have analyzed the grain data and there is also *positive* correlation across genotypes between grain yield in the late-season stress treatment and early-season stomatal conductance (or whole-plant transpiration). We also note that the biomass data reflect only above-ground biomass, and as such, should not be interpreted as a whole-plant growth.

**Fig. 3.**
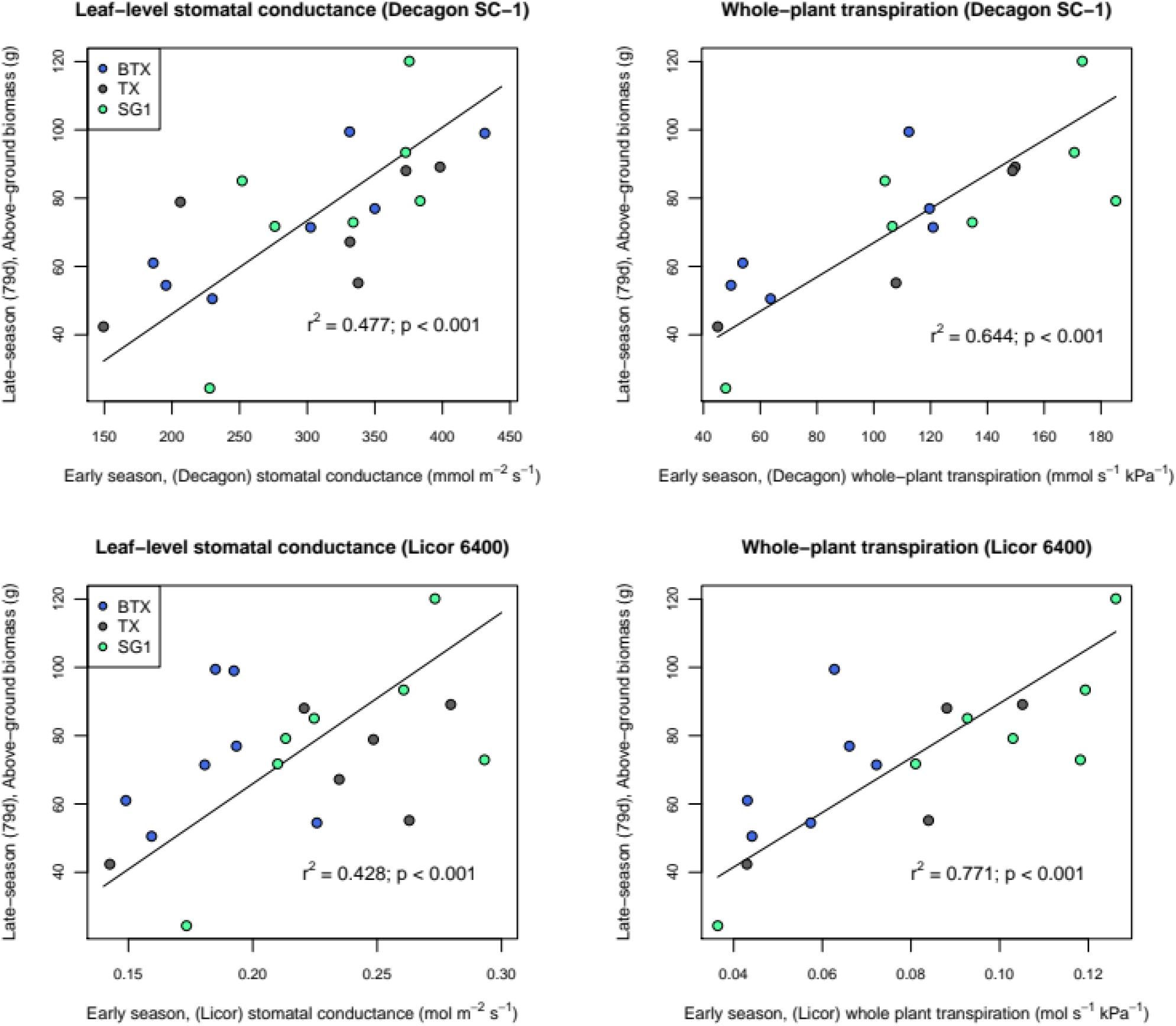
Relationship between early-season stomatal conductance (x axis) and late-season above-ground biomass (y axis) measured using the Decagon SC-1 steady-state porometer (top two panels) and the Licor-6400 (bottom two panels), and expressed at the leaf-level (left two panels) or whole-plant level (right two panels), scaled up using leaf area data collected on day 8/27/2022. Symbols represent mean genotype plot values.

## Discussion

The strong and well-understood physiological linkage between water use and carbon gain presents an important obstacle for limited transpiration theory and the practical application of this theory. For limited transpiration traits to confer improved performance, the carbon lost by limiting water use during the early season must be overcome by improved growth and plant water status (i.e., water potential) later in the season. This hinges critically on assumption that water can be stored in soil at the same plant-available water potential for weeks/months, without being lost to competing sinks (evaporation, unsaturated flow to drier soil layers, and use by weeds) (Tanner and Sinclair 1983; Sinclair 2018). This assumptions appears questionable and we have not been able to find a convincing study that demonstrates this principal, and should be even more unlikely in arid/semi-arid climates where soil and atmospheric aridity are high, as well as for shallow-rooted species. In support of our skepticism on this point, soil water storage efficiency has been reported as low as 30% in the Australian wheat belt (Sprigg et al. 2014). However, considering this, and given that this study was conducted under only high VPD conditions, it is possible that the water use and above-ground growth patterns (e.g., positive correlation between early-season water use and above-ground growth) may have weakened, or even exhibited inverse correlation, if this study had been conducted in a more humid environment with deep antecedent soil water present.

Even if soil water can indeed be transferred without meaningful loss from early season to late season, how should we assign an opportunity cost to water that is *not* spent early in the crop’s development? Water that is spent early in the season can be invested in root growth that would allow for improved access to deep soil water later in the season. As such, limited transpiration crops would appear to be purposely ill suited to access “saved” deep soil water by nature of their design. Although we think it is a fairly safe assumption that this early-season carbon would be important for resource capture and reproductive development, we did not measure allocation patterns in our study (e.g., no root or non-structural carbohydrates measurements), and therefore do not know what the opportunity costs of poor early-season growth (lower water use) might be.

In our view, breeding plants to use less water at any point of their development disregards a well-aligned physiological system that has been designed by natural and artificial selection to convert water into carbon. It is also logical, in our view, given the value of early-season carbon to a vegetative plant, that carbon acquired early in the season cannot be “traded” for carbon (or water) later in the season without very significant opportunity costs, although we think this remains an important research question. A breeding strategy that focuses solely on providing higher water status during grain development, and gives short shrift to the physiological and developmental consequences of reduced carbon income during the vegetative phase, is almost certainly not the best strategy for improving plant growth in arid/sub-arid environments. Rather, we advocate for a strategy that improves access to deep soil water (deeper root systems, higher fine root density), the near-complete extraction of this water (low operating water potential), as well as aligned xylem transport, photosynthesis, sugar export (source→sink), and stomatal traits that *maximize* carbon gain (not grain yield per se). We note that this strategy places a premium on maximizing total water extraction and use, rather than improving water use efficiency (aka: transpiration efficiency), reducing transpiration, or shifting soil water from early season to late season use (aka: “effective” water use).

